# S1P-S1PR1 activity controls VEGF-A signaling during lymphatic vessel development

**DOI:** 10.1101/845396

**Authors:** AM Golding-Ochsenbein, S Vidal, B Wilmering Wetter, C Guibourdenche, C Beerli, L Chang, S Leonhard, N Holway, K Seuwen, G Jurisic

## Abstract

Sphingosine-1-phosphate (S1P), a lipid signaling molecule produced by endothelial cells, is required for development and homeostasis of blood vessels. However, its role during lymphatic vessel development is unclear. We show in murine newborns that pharmacologically enhanced S1P signaling increases VEGF-A-dependent LEC proliferation. In contrast, S1PR1 inhibition, mediated by the antagonist NIBR0213 or LEC-specific genetic deletion of *S1pr1*, promotes filopodia formation and vessel branching, independent of VEGF-A. To investigate the S1P and VEGF-A signaling crosstalk observed *in vivo*, we used LECs cultured *in vitro.* We demonstrate that S1P activates endogenous S1PR1 in a constitutive, autocrine manner. Importantly, S1P-S1PR1 activity was required for VEGF-A-induced LEC proliferation and strongly supported ERK1/2 activation and VEGFR-2 trafficking to the perinuclear area. In conclusion, S1P-S1PR1 signaling promotes VEGF-A-dependent LEC proliferation and limits migratory and filopodia-forming responses. Hence, S1P-S1PR1 signaling is required for balanced growth factor-induced lymphangiogenesis and correctly patterned lymphatic vessels during postnatal development.

## Introduction

Lymphangiogenesis, the formation of new lymphatic vessels from pre-existing ones, is orchestrated by tissue-derived growth factors such as VEGF-C and VEGF-A, and by cell intrinsic mechanisms (Vaahtomeri et al., 2017) that create a balance between proliferating and migratory, filopodia-forming lymphatic endothelial cells (LECs) (James et al., 2013). This balance is key to subsequent vascular patterning events such as branching and hence, influence lymphatic vessel morphology (Gerhardt et al., 2003). Interestingly, VEGF-A is known to enhance proliferation and less so sprouting (Nagy et al., 2002; Wirzenius et al., 2007), whereas VEGF-C induces both sprouting and proliferation.

VEGF-A activates the receptor VEGFR-2, whereas VEGF-C, in its processed form, activates both VEGFR-2 and VEGFR-3. The latter receptor is critically required for lymphangiogenesis. Both receptors belong to the receptor tyrosine kinase family, and ligand activation leads to dimerization and auto-phosphorylation, followed by internalization (reviewed in (Zhang and Simons, 2014)) and fusion with endosomal compartments (Simons et al., 2016; Zhang and Simons, 2014). These events are important for the activation of several downstream signaling pathways including ERK1/2 that is crucial for endothelial cell function (Simons et al., 2016).

Sphingosine-1-phosphate (S1P), a lipid mediator secreted by erythrocytes, platelets and endothelial cells, signals via 5 specific G protein-coupled receptors (S1PR1-5) by inducing their internalization and activating downstream signaling cascades (Mullershausen et al., 2009; Yanagida and Hla, 2017). S1PR1, highly expressed by endothelial cells, is crucial for embryonic blood vessel development and function (Yanagida and Hla, 2017). It stabilizes endothelial barrier and negatively regulates blood endothelial cell sprouting (Ben Shoham et al., 2012; Bigaud et al., 2016; Gaengel et al., 2012; Jung et al., 2012). However, opposing conclusions of S1P acting either pro- or anti-angiogenic were previously made (Ben Shoham et al., 2012; Caballero et al., 2009; English et al., 2001; English et al., 2000; Gaengel et al., 2012; Licht et al., 2003). For lymphatic vessels, S1P was reported to promote lymphangiogenesis in a matrigel plug assay in adult mice (Yoon et al., 2008). Additionally, sphingosine kinase activity in LECs was shown to be required for generation of S1P and proper lymphatic vessel patterning and arrangement of cell junctions *in vivo* (Pham et al., 2010). However, the mechanism of how S1P affects lymphatic vessel development, in particular its effect on growth factor signaling, and the role of S1PR1 during this process are not known.

We show here, for the first time, that S1P-S1PR1 activity influences lymphangiogenesis by promoting VEGF-A-dependent LEC proliferation and by limiting migratory and filopo-dia-forming responses. Furthermore, antagonism of S1PR1 leads to dysfunctional lymphatic vessels. Hence, S1P-S1PR1 signaling is required for the correct patterning and maintenance of function of the lymphatic vessel network during postnatal development.

## Methods

### Endothelial cell culture

Primary adult human dermal LECs (PromoCell) or human umbilical vein endothelial cells (HUVECs) (PromoCell) were cultured on plates coated with 50 μg/ml Purecol collagen type I (Advanced Biomartix) and in EGM-MV medium without VEGF-A (PromoCell). For starvation of the cells, EBM-MV (PromoCell) without FCS was used. For washing steps pH7.4 phosphate-buffered saline (PBS) without CaCl_2_ and MgCl_2_ (Gibco) was used. Cells were treated with the dual non ATP-competitive sphingosine kinase inhibitor SKI-II (SIGMA), S1P (Avanti Polar) diluted in 90% DMSO containing 50 mM HCl, NIBR0213 (Novartis) and recombinant carrier-free human VEGF-A165 (R&D systems).

### Measurements of S1P concentration

S1P concentrations were measured as previously described (Billich et al., 2013). To reduce unspecific binding, extracts were subjected to acetylation as described in (Berdyshev et al., 2005). Sphingosine derivatives were detected in positive mode; negative ionization was used for S1P. Quantification was performed based on the area ratios of the compound over internal standard in the extracted ion chromatograms. Recovery was >86% for all analytes. The limit of quantification, as determined by the lowest calibration sample showing signal-to-noise ratio >5 and accuracy <14%, was 0.5 ng/ml for Sph, 0.1 ng/ml for S1P.

### Electric Cell-substrate Impedance Sensing (ECIS)

The lymphatic endothelial cell barrier tightness was assessed using an ECIS® ZΘ instrument with a 16-well array station (Applied Biophysics). 8-well chamber slides (8W10E+, ibidi) were pre-incubated with 50 μg/ml fibronectin (Roche) and then washed with sterile H_2_O. Next, 30000 adult human dermal LECs per well were plated in EGM and impedance values were recorded at 4,000 Hz every 30 seconds. When a monolayer with stable impedance levels of 3000-3500 ohm was formed, treatments with compounds started. To investigate LEC endogenous S1P functions, 10 μM SKI-II or DMSO control was spiked into corresponding wells. After 4 h cells were washed with PBS and starved in EBM containing again 10 μM SKI-II for 3 h. Next, cells were treated ± 1 μM S1P, S1P+0.3 μM NIBR0213 or DMSO control. To rule out NIBR0213 toxic effects, 0.1 or 0.01 μM NIBR0213 was spiked into corresponding wells. After 5 h 10 μM S1P or DMSO control were spiked into corresponding wells. Graphs were generated using TIBCO Spotfire 6.5.3. Impedance curves were normalized to DMSO control.

### Immunofluorescent approaches for in vitro experiments

Cells were starved for 4 h or overnight in medium without FCS and received indicated treatments for indicated times. Then cells were fixed with 4% PFA for 10 min at room temperature and washed with PBS before blocking for 1 h in immunomix (PBS containing 5 % donkey serum, 1 % bovine serum albumin (BSA) and 0.1 % Triton-X). The following antibodies were incubated overnight at 4°C: rat anti-human CD31 (DAKO, M082329-2), rabbit anti-human S1PR1 (Santa Cruz, sc-25489), goat anti-human VEGFR-2 (R&D systems, AF357), rabbit anti-human Rab5 (Cell Signaling, #3547P). After an intense wash step in 0.1% Triton-X PBS, samples were incubated for 1 h at room temperature with Alexa Fluor 488, 594 or 647 nm-conjugated secondary antibodies (Invitrogen). Samples were imaged with a Zeiss LSM 700 (Zeiss) confocal microscope using the Zen 2011 SP3 (black edition) software or for VEGFR-2 trafficking with a Yokogawa CV7000 system using the CellVoyager R1.17.05 software. Yokogawa images were analyzed with ImageJ: The perinuclear area was defined by Hoechst area plus a 1.625 μm radius and total cell area was defined based on CD31 plasma membrane staining. Total cell VEGFR-2 was defined as 100%. Values were normalized to DMSO control.

### Homogeneous time resolved fluorescence (HTRF)

HTRF approaches (Cisbio) were used to determine VEGFR-2 pTyr1175 and ERK pThr202/Tyr204 levels. To measure VEGFR-2 pTyr1175 levels after VEGF-A stimulation with or without S1PR1 pathway modulation, 40000 cells per 96 well were seeded in EGM in the morning. In the evening, cells were washed with PBS and starved overnight in EBM without FCS. Next, cells were stimulated with the indicated treatments for 2 and 5 min and 2 h before processing according to manufacturer’s instructions. For determination of pERK1/2 levels in the presence or absence of SKI-II, 20000 human dermal LECs per 96 well were seeded in EGM. The next day 10 μM SKI-II was spiked into corresponding wells and incubated for 4 h before EGM removal, a PBS wash and the addition of EBM starvation medium without FCS ± 10 μM SKI-II. Cells were starved for 3 h before 10 min stimulation with the indicated treatments. Next, cells were processed according to manufacturer’s instructions. Fluorescent signals were measured with TRF Light Unit EnVision(PerkinElmer). EC_50_ and E_max_ were calculated with Prism 6 (GraphPad Software).

### Cell proliferation assay

For the 5-ethynyl-2’-deoxyuridine (EdU) proliferation assay, 5000 LEC per well of a 96 well plate were seeded in EGM in the morning. In the evening, cells were washed with PBS before an overnight starvation in EBM. The next morning, cells were treated with 100 ng/ml VEGF-A ± 0.3 μM NIBR0213 or DMSO control and in the evening 10 μM EdU was spiked. 24 h after the addition of VEGF-A cells were fixed with 4 % PFA before further processing according to the Click-iT EdU Imaging Kit (Invitrogen). Experiments were performed with eight replicates per group. Per well nine non-overlapping images were acquired and analyzed with the Operetta High Content Screening System microscope (PerkinElmer) and Harmony 4.1 software.

### Mice

For *in vivo* experiments with pharmacological approaches C57BL/6 females were purchased from Charles River Laboratories (France) or came from Novartis Pharma AG Basel internal breeding facilities. For lymphatic vessel-specific depletion of S1PR1, the well-established mouse lines Prox-1creERT2 mice (Bazigou et al., 2011) and S1PR1 floxed mice (Allende et al., 2003) were crossed and bred in the animal breeding facility of Novartis Pharma AG, Basel. For pharmacological approaches C57BL/6 pups received i.p. 4 or 20 mg/kg/day Compound 31 (Cmpd 31) (Novartis) diluted in PEG200/5% glucose (70/30%) or PBS/PEG200/5% glucose (50%/35%/15%) for co-treatments, 3 or 6 mg/kg/day VEGF-A neutralizing antibody 4G3 (Novartis) diluted in PBS/PEG200/5% glucose (50%/35%/15%), 1, 5 or 10 mg/kg/day NIBR0213 (Novartis) diluted in PEG200/5% glucose (70/30%) or PBS/PEG200/5% glucose (50%/35%/15%) for co-treatments with 4G3 (6 mg/kg/day) from postnatal day (P) 0.5 to P3.5 and were sacrificed at P4.5. To induce recombination events, homozygous *S1pr1* flx and *Prox-1-CreERT2+* (S1PR1iΔLEC) or *Prox-1-CreERT2-* (control) littermates received intragastric injections of 50 μg tamoxifen (Sigma), dissolved in EtOH and sunflower seed oil (Sigma) from P1 to P4. Tissues were harvested at P7 to allow enough time for recombination to occur.

### Animal study approval

All *in vivo* experiments were approved by the Cantonal Veterinary Office Basel-Stadt (license number BS-2586).

### Whole-mount staining and morphometric analyses

Abdominal skin or diaphragms were harvested and skin samples were incubated for 1 h in 20 mM EDTA at 37°C to facilitate the removal of the epidermis. Then tissues were fixed for 2 h in freshly prepared 4% PFA at 4°C. Next, samples were washed in PBS and blocked in immunomix (described above) for 1 h at RT before overnight primary antibody incubation at RT. The following primary antibodies were used: rat-anti mouse S1PR1 (R&D Systems, MAB7089), hamster anti-mouse CD31 (Millipore, MAB1398Z), goat antimouse Prox-1 (R&D Systems, AF2727), rabbit anti-mouse LYVE1 (AngioBio, 11-034), goat anti-mouse LYVE1 (R&D Systems, AF2125), rabbit anti-mouse KI-67 (Neomarkers, RM-9106-S1), goat anti-mouse VEGFR-2 (R&D Systems, AF644). The next day, whole mounts were washed and incubated for 2 hours with AlexaFluor 488, 594 or 647-conjugated secondary antibodies (all from Invitrogen) at RT. Whole-mount z-stacks were acquired on a Zeiss LSM 700 confocal microscope with the Zen 2011 SP3 (black edition) software. Images were processed with Zen software. For lateral diaphragm segments images were processed with a personalized Fiji macro (Schindelin et al., 2012) (Rueden et al., 2017). The macro automated the following tasks: the maximum intensity projection of each image stack was calculated and then each projection was stitched together using Fiji’s Grid/Collection stitching plugin (Preibisch et al., 2009). The stitched image was then thresholded using Fiji’s default (IsoData) algorithm (Ridler and Calvard, 1978) and automatic threshold value, colored green and then saved. The macro is freely available at https://github.com/Novartis/ExactAddressTBC. Lateral diaphragm segments were analyzed as described previously (Ochsenbein et al., 2016). Other images were analyzed by hand with ImageJ.

### Quantification of lymphangiogenic growth factor levels in P2.5 diaphragms

To assess growth factor levels in postnatal diaphragms, tissues were collected and mRNA levels were assessed by qPCR. To do so, two diaphragms of P2.5 C57Bl6 pups were collected in RNA later (Qiagen) before they were processed in RLT buffer containing 1% mercaptoethanol (Sigma) with the help of metal beads (MP Biomedicals) and the FastPrep-24 tissue ruptor (MP Biomedicals). Next, RNA was extracted according to the RNeasy Micro Kit (Qiagen). The RNA was transcribed to cDNA using the High-Capacity cDNA Reverse Transcription Kit (Applied Biosystems). The PCR reactions were performed using a Quant Studio 7 Flex Fast Real-Time PCR System (Applied Biosystems by life technologies) and the following Taqman primers (Thermo Fisher): *Actb* (Mm02619580_g1), *Vegf-A* (Mm00437306_m1), *Vegf-C* (Mm00437310_m1), *Vegf-D* (Mm01131929_m1). Data is presented by fold change to *Actb*.

### Statistical data analysis

Statistical analyses were performed using Prism 7 (GraphPad Software). Data are shown as mean ± standard deviation. Normal distribution was tested with Shapiro-Wilk test. Normally distributed data were analyzed using the student’s t-test or, for comparisons of means of more groups, one-way ANOVA and Dunnett’s test were used. For non-normal distributed data non-parametric tests where used. Statistical significance is indicated by asterisks: * p < 0.05, ** p < 0.01, *** p < 0.001, **** p< 0.0001. Differences were considered statistically significant when p < 0.05.

## Results

### Increasing tissue S1P levels enhances lymphatic endothelial cell proliferation in vivo in a VEGF-A-dependent manner

To study the role of S1P in early postnatal lymphangiogenesis, we used a previously established murine model where lymphatic vessels on the pleural side of the diaphragmatic muscle are monitored (Ochsenbein et al., 2016). In a first series of experiments we enhanced S1P signaling by treating pups from postnatal day (P) 0.5-3.5 with Cmpd 31, a highly specific S1P lyase inhibitor that increases S1P concentrations *in vivo* (Weiler et al., 2014). At P4.5, LYVE1+ lymphatic vessels expressed S1PR1, which was also present on some single cells and on CD31+ blood vessels (Fig. 1A). Intraperitoneal injection of 20 mg/kg/day of Cmpd 31 significantly increased S1P levels in muscle tissue (Fig. S1A) and, in the diaphragm, lymphatic vessel diameters and lymphatic endothelial cell (LEC) proliferation (assessed as %KI-67+ cells per PROX-1+ cells) without changing the vessel branching response (Fig 1B-E,G,H). Thus, we conclude that enhanced S1P signaling boosts LEC proliferation during postnatal development.

**Figure 1:**
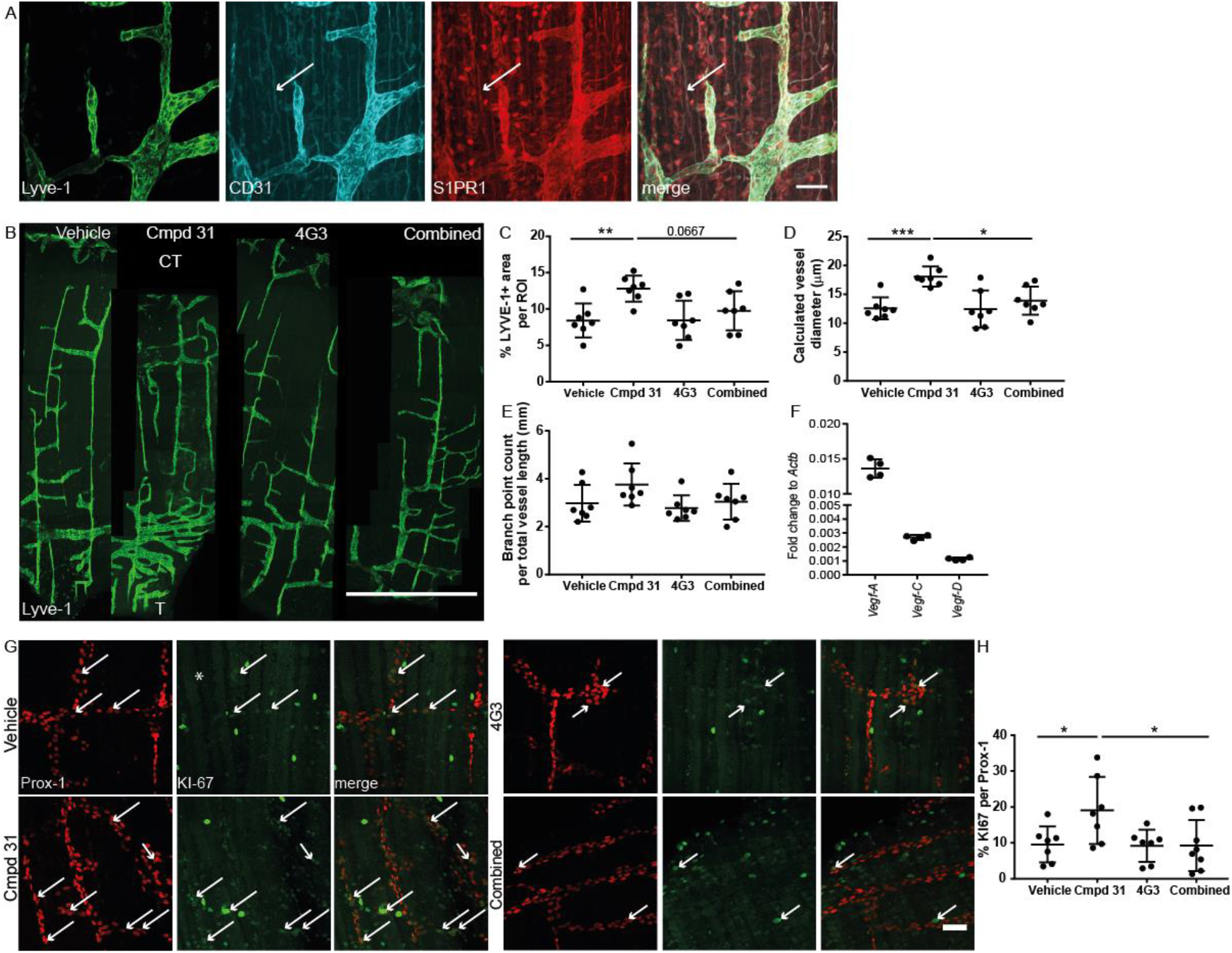
S1P enhances LEC proliferation in a VEGF-A-dependent manner *in vivo*. A) Representative confocal images of P4.5 diaphragm whole mounts (pleural side) stained for LYVE1 (green), CD31 (blue) and S1PR1 (red). Arrow highlights an example of CD31+ and S1PR1+ blood vessel. Scale bar 50 μm. B) Representative stitched confocal images of P4.5 LYVE1-immunostained diaphragm whole mounts (pleural side) of pups exposed to indicated treatments. T= Thorax wall, CT = Central tendon, scale bar 1 mm. C–E) Quantification of lymphatic vessel parameters of treated pups. F) Real-time qPCR analysis of lymphatic growth factor gene expressions compared to *Actb* in P2.5 diaphragms. G-H) Representative confocal images and quantifications of diaphragm whole mounts immunostained for Prox-1 (red) and KI-67 (green) after indicated treatments. Arrows: KI-67 + LECs, scale bar 50 μm. Statistical analyses: Dots represent animal mean, lines represent group means, error bars represent ±SD. Statistical significance was calculated with one-way ANOVA and Dunnett’s test.

Next, we asked whether S1P might increase lymphatic vessel diameter and cell proliferation via cross-talk with VEGFR-2, as reported for blood endothelial cells (Gaengel et al., 2012). Indeed, VEGFR-2 is expressed on LYVE-1 positive lymphatic vessels at P2.5 (Fig S1B) and *Vegf-a* is the most abundantly expressed endothelial growth factor in the postnatal diaphragm at P2.5 (Fig 1F). We therefore treated neonates from P0.5-3.5 with Cmpd 31 (20 mg/kg/day i.p.) with or without 3 mg/kg/day of 4G3, a VEGF-A neutralizing antibody (Poor et al., 2014). The analysis of P4.5 LYVE1 immuno-stained diaphragms revealed that VEGF-A neutralization alone had no detectable effect on lymphatic vessel parameters in the present model, as expected from earlier experiments (Zhang et al., 2018). However, VEGF-A neutralization mildly reduced the Cmpd 31-mediated increase in LYVE1+ lymphatic vessel area and entirely abolished both the Cmpd 31-induced enlargement of lymphatic vessel diameter and LEC proliferation (Fig 1B-E,G,H). None of the tested treatments significantly affected vessel branch formation. Cmpd 31 also increased lymphatic vessel area in the abdominal skin with a trend of reduction by co-treatment with 4G3 (Fig S1C). Hence, we report that enhanced S1P signaling enlarges lymphatic vessel diameters and boosts LEC proliferation in a VEGF-A-dependent manner *in vivo*, without affecting vessel branching.

### Lymphatic endothelial cell S1PR1 signaling reduces lymphatic vessel sprouting independently of VEGF-A

To complement our pharmacological approach of investigating S1P signaling, we used the specific S1PR1 competitive antagonist NIBR0213 (Quancard et al., 2012) during early postnatal lymphangiogenesis. We treated pups from P0.5 to P3.5 i.p. with 10 mg/kg/day of NIBR0213 and sacrificed them at P4.5. At this time point, all NIBR0213 -treated pups had developed chylus ascites, indicative of malfunctioning lymphatic vessels (Fig 2A). Interestingly, analyses of diaphragm whole mounts showed that inhibition of S1PR1 *in vivo* only mildly reduced LEC proliferation and had no effect on LYVE1-positive lymphatic vessel area, the total lymphatic vessel network expansion or the calculated lymphatic vessel diameter (Fig 2B, Fig S2A). However, NIBR0213 led to a strong, dose-dependent increase in branch formation (Fig 2B, Fig S2B) and enhanced filopodia formation (Fig 2B, Fig S2C), suggesting that S1PR1 activity inhibits LEC migratory, filopodia-forming responses *in vivo*. Neutralization of VEGF-A did not alter NIBR0213-mediated enhanced branch formation (Fig S2D, E).

**Figure 2:**
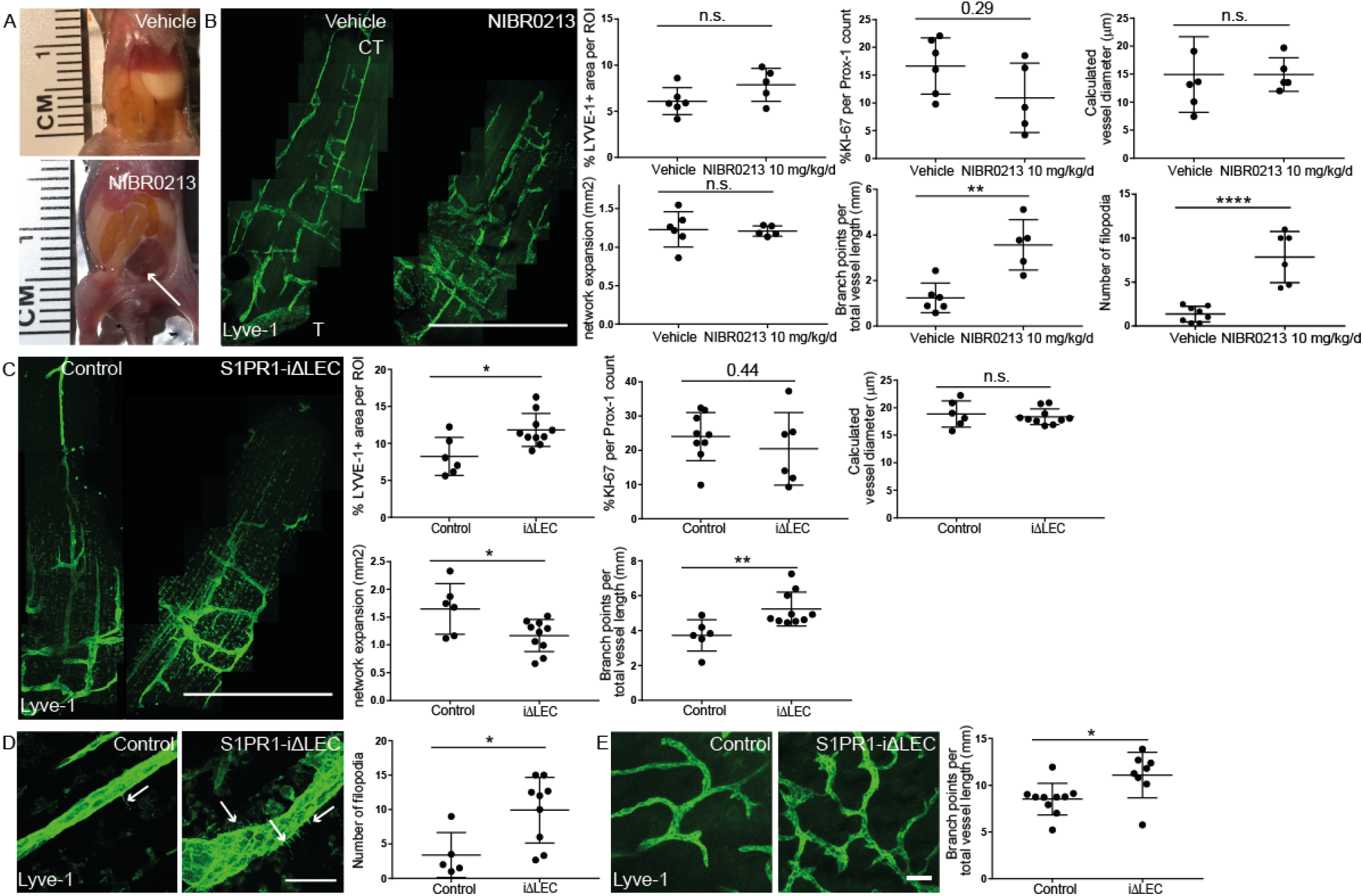
S1PR1 in lymphatic endothelial cells reduces filopodia formation and branching. A) Images of P4.5 vehicle and NIBR0213 treated pups with exposed peritoneum. Arrow indicates chylus ascites. B) Stitched confocal images and quantification of lateral segments of LYVE1 immunostained diaphragm whole mounts of vehicle or 10 mg/kg/day NIBR0213 treated pups. CT= Central tendon, T = Thorax wall. Scale bar 1 mm. C) Stitched confocal images and quantification of a lateral segment of LYVE1 immunostained diaphragm whole mounts of P7 control or S1PR1-iΔLEC pups. Scale bar 1 mm. D) High magnification confocal images and quantification of increased filopodia formation of LYVE1 immunostained diaphragmatic lymphatic vessel in control and S1PR1iΔLEC pups. Arrow indicates filopodia, scale bar 50 μm. E) Confocal images and quantification of LYVE1 immunostained abdominal skin whole mount of P7 control or S1PR1iΔLEC pups. Scale bar 50 μm. Statistical analysis: Dots represent animal mean, lines represent group means, error bars = ±SD. Statistical significance was calculated using an unpaired Student’s t-test, or for C Branch points per vessel length a Mann Whitney test.

To demonstrate that the effects of S1PR1 antagonism are indeed intrinsic to LECs we used an inducible genetic loss-of-function approach to delete *S1pr1* specifically in this cell type. Homozygous *S1pr1* flx and *Prox-1-CreERT2+* or *Prox-1-CreERT2−* littermates were treated intragastrically from P1-4 with 50 μg/day tamoxifen and diaphragms were harvested at P7. Analyses of diaphragm whole mounts revealed that, compared to homozygous *S1pr1* flx and *Prox-1-CreERT2−* control pups, S1PR1 was strongly decreased on LYVE1+ lymphatic vessels in homozygous *S1pr1* flx and *Prox-1-CreERT2+* (S1PR1iΔLEC) littermates, while the S1PR1 expression on blood vessels remained intact (Fig S2F). Importantly, the lymphatic endothelial cell-specific S1PR1 loss-of-function recapitulated the phenotypes caused by NIBR0213 treatment. Lymphatic vessel branching and filopodia formation were strongly increased whereas no changes in lymphatic vessel diameter and LEC proliferation were found (Fig 2C, D, Fig S2G). Deficiency of LEC S1PR1 led to a decreased lymphatic network expansion, suggesting that lymphatic vessel growth was inefficient (Fig 2C). Superficial lymphatic vasculature in the dermis likewise showed increased branch formation, implying a systemic and not an organ-specific effect (Fig 2E). Thus, LEC-specific S1PR1 signaling strongly inhibits lymphatic endothelial cell filopodia formation and vessel branching and is required for a balanced expansion and proper function of the lymphatic vasculature during postnatal development.

### Lymphatic endothelial cells in vitro exhibit autocrine and constitutive S1P-S1PR1 signaling

To study the molecular mechanism by which S1P regulates lymphatic vessel development, we characterized S1P production and signaling in primary human dermal lymphatic endothelial cells *in vitro*. Liquid chromatography–mass spectrometry analyses of LECconditioned medium showed that LECs secrete high amounts of S1P (Fig 3A). Treatment with SKI-II, an inhibitor of sphingosine kinases 1+2 that phosphorylate sphingosine to S1P, completely abolished S1P production. Of note, we showed that already plain, unconditioned commercial endothelial growth medium (PromoCell EGM-MV) contains ca. 6 nM of S1P (Fig 3A) and conclude that active concentration of S1P is constantly present in regular LEC *in vitro* culture.

**Fig 3:**
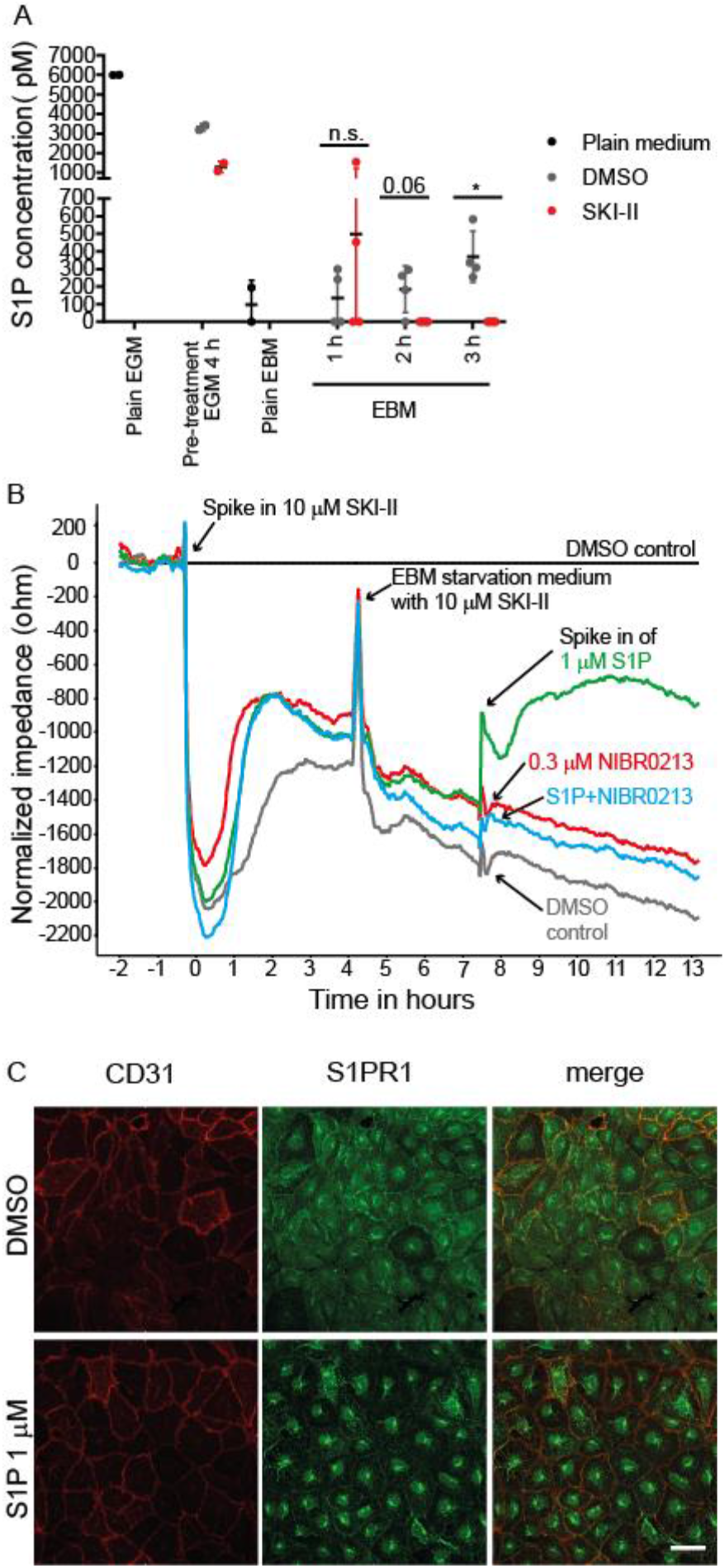
Lymphatic endothelial cells *in vitro* exhibit autocrine and constitutive S1P-S1PR1 signaling. A) S1P concentrations in plain medium (black dots) or in conditioned supernatants of LEC treated ± SKI-II (grey or red dots). B) Normalized (to DMSO control) electrical impedance of LEC monolayers after indicated treatments. C) CD31 (red) and S1PR1 (green) immunofluorescence of LECs monolayer treated with 1 μM S1P for 1 h. Representative confocal images; Scale bar = 50 μm. Statistical analyses: Dots represent individual measurements, lines group means, error bars = ±SD. Statistical significance was calculated using an unpaired Student’s t-test and Welch’s correction for unequal variances.

Next, we assessed whether S1P can act on LECs directly in an autocrine manner by measuring endothelial barrier function, a process known to be modulated by S1P (Xiong and Hla, 2014), using electric cell-substrate impedance sensing (ECIS) with high temporal resolution. Inhibition of autocrine S1P production by SKI-II treatment led to the destabilization of the LEC barrier and this effect could be overcome by the addition of exogenous S1P (Fig 3B). We then used the competitive and selective antagonist NIBR0213 and showed that S1P-induced barrier stabilization was fully reversed, demonstrating the involvement of S1PR1. NIBR0213 is not toxic for LECs as excess S1P could compete for S1PR1 binding and overcome NIBR0213-induced barrier opening (Fig S3). Thus, LECs produce S1P that acts via S1PR1 in an autocrine and constitutive signaling loop that manifests in LEC barrier function regulation. Furthermore, LECs expressed S1PR1 protein that was internalized upon stimulation with a saturating concentration of 1 μM exogenous S1P (Fig 3C).

### The mitogenic effect of VEGF-A on lymphatic endothelial cells in vitro depends on S1P-S1PR1 signaling

In order to gain a better mechanistic understanding of the interaction between VEGF-A and S1P signaling resulting in the enlarged lymphatic vessel diameters and LEC proliferation *in vivo*, we studied VEGF-A-mediated proliferation of human dermal LECs *in* vitro in the presence or absence of S1PR1 signaling. VEGF-A at 100 ng/ml significantly increased cell proliferation as determined by EdU incorporation, and strikingly, this effect was completely inhibited by NIBR0213 (Fig 4A). Thus, S1PR1 signaling is required for VEGF-A-mediated LEC proliferation *in vitro*.

**Figure 4:**
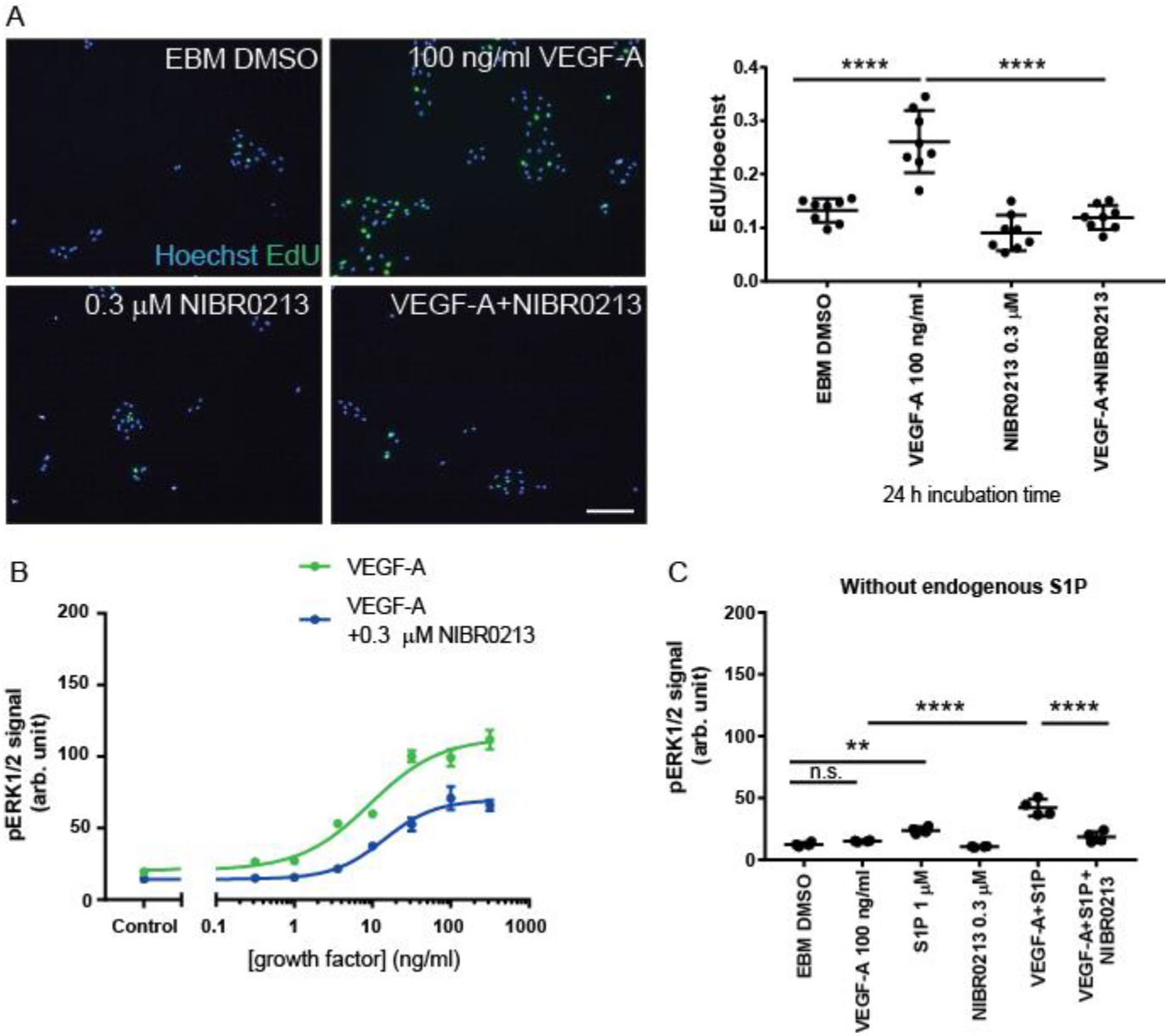
VEGF-A-induced ERK1/2 activation and proliferation depend on autocrine S1P-S1PR1. A) EdU (green) and Hoechst (blue) staining along with quantifications of cultured LEC after 24 h of indicated treatments. Representative images; Scale bar: 200 μm. B,C) Quantification of pERK1/2 levels assessed by HTRF in LEC lysates after 10 min of indicated treatments. Statistical analyses: In A and C, dots represent individual measurements, lines represent group means. In B, dots represent group means, lines represent fitted curves. Error bars = ±SD. Statistical significance was calculated using one-way ANOVA and Dunnett’s test.

### Autocrine S1P-S1PR1 signaling controls VEGF-A-induced ERK1/2 activation

We went on to study growth factor-induced ERK1/2 activation *in vitro*, a highly relevant signaling node for mitogenesis in general (Simons et al., 2016) and for VEGF-A down-stream activity in particular (Wang et al., 2010). Using a high throughput homogeneous time resolved fluorescence (HTRF) approach, we determined the phospho-ERK1/2 (pERK1/2) response after 10 minutes of titrated VEGF-A with and without NIBR0213 (Fig 4B). Indeed, NIBR0213 inhibited pERK1/2 activation mediated by VEGF-A and caused a significant increase in EC_50_ of VEGF-A and strongly decreased the Emax of VEGF-A down to 50% (Table 1). Thus, we conclude that VEGF-A-induced ERK1/2 activation is strongly dependent on S1PR1 signaling. Furthermore, at 10ng/ml VEGF-A, a sub-saturating concentration, the addition of exogenous S1P significantly increased growth factorinduced ERK1/2 activation (Fig S4A). Hence, S1P-S1PR1 signaling enhances and S1PR1 antagonism limits VEGF-A-mediated ERK1/2 activation in LECs *in vitro*.

**Table 1.**
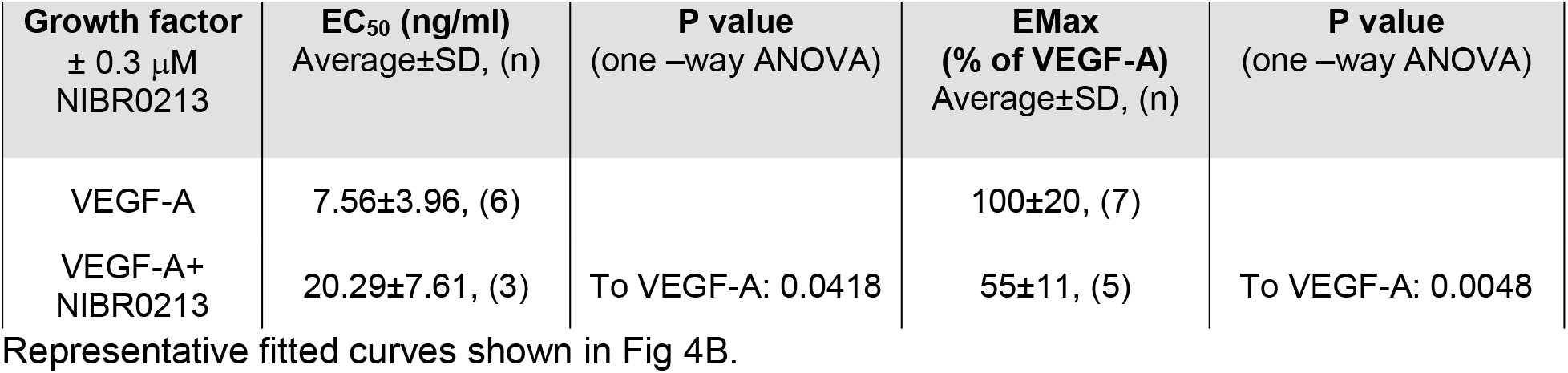
Potency and efficacy parameters for VEGF-A induced ERK1/2 activation.

A positive signaling cross-talk between S1PR1 and VEGFR-2 in LECs was unexpected as earlier work in HUVECs suggested a decrease of VEGF-A-induced pERK1/2 levels (Gaengel et al., 2012). We therefore carried out experiments with human umbilical vein endothelial cells (HUVECs) and obtained results similar to the ones reported here for LECs. Addition of exogenous S1P 1 h prior to or simultaneously with VEGF-A increased pERK1/2 levels and NIBR0213 significantly reduced the signal (Fig S4B).

To further confirm that growth factor-induced ERK1/2 activation depends on autocrine production of S1P we assessed the pERK1/2 response after VEGF-A treatments in the presence of the sphingosine kinase inhibitor SKI-II (Fig 4C). With SKI-II, VEGF-A was not able to induce ERK1/2 activation significantly compared to control. Co-stimulation with exogenous S1P overcame the SKI-II block for VEGF-A and significantly increased pERK1/2 levels compared to VEGF-A alone (Fig 4C). This increase was inhibited by S1PR1 antagonism.

Taken together, these results show that VEGF-A-induced ERK1/2 activation strongly depends on autocrine S1P-S1PR1 signaling.

### VEGF-A-induced VEGFR-2 trafficking to the perinuclear area depends on S1P-S1PR1 activity

To investigate how S1P-S1PR1 affects VEGF-A-mediated signaling events, we studied VEGFR-2 phosphorylation and intracellular trafficking following VEGF-A treatment with or without pharmacological S1P pathway modulation. These events, together with receptor internalization, were described to be critical for growth factor action (Simons et al., 2016; Wang et al., 2010). Similarly, S1PR1 receptor internalization, trafficking to perinuclear area and persistent signaling from intracellular compartments, were demonstrated to be relevant for endothelial barrier regulation (Mullershausen et al., 2009). As shown in Fig. 5A, VEGF-A significantly increased VEGFR-2 Tyr1175 phosphorylation that was unaffected by the S1PR1 antagonist NIBR0213 at all tested time points and VEGFA concentrations. This indicates that S1P-S1PR1 is not involved in the initial steps of VEGFR-2 signaling, i.e. receptor dimer formation and autophosphorylation. We next assessed whether NIBR0213 co-treatment affected VEGFR-2 co-localization with Rab5, an early endosome marker. The co-localization was similar with NIBR0213 co-treatment compared to VEGF-A alone at all tested time points (Fig S5). However, further onward VEGFR-2 trafficking to the perinuclear region was significantly reduced by NIBR0213 cotreatment compared to VEGF-A alone after 10, 20 and 30 min of treatment (Fig 5B, C). These data show that S1P-S1PR1 signaling is not required for VEGFR-2 trafficking to the early endosome but for VEGF-A-mediated VEGFR-2 trafficking to the perinuclear region.

**Figure 5:**
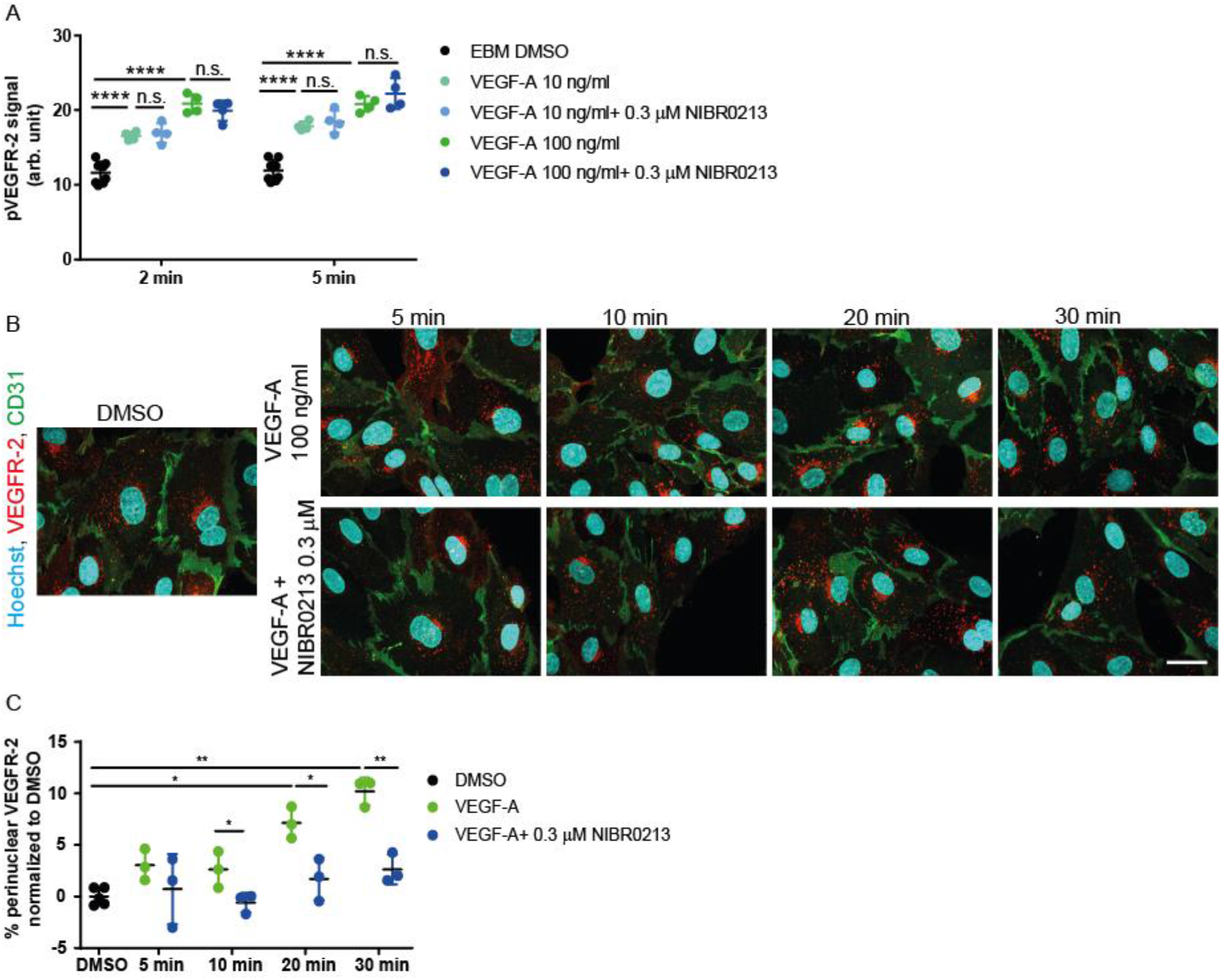
VEGF-A-induced VEGFR-2 translocation to the perinuclear area depends on S1PR1. A) Quantification of pVEGFR-2 levels, assessed by HTRF, in LEC lysates after indicated treatments and incubation times. B,C) Immunofluorescent staining of VEGFR-2 (red), CD31 (green) and Hoechst (blue) and quantification of VEGFR-2 trafficking to the perinuclear area of cultured LEC after indicated treatments and incubation times. Representative confocal images; Scale bars: 50 μm. Statistical analyses: Dots represent individual measurements, lines group means, error bars = ±SD. Statistical significance was calculated using one-way ANOVA and Dunnett’s test.

## Discussion

Studies have demonstrated that S1P-S1PR1 signaling is required for development and homeostasis of blood vessels (Ben Shoham et al., 2012; Gaengel et al., 2012; Jung et al., 2012). In addition, a signaling cross talk between VEGFR-2 and S1PR1 was reported in blood endothelial cells (Gaengel et al., 2012). However, in the past different groups have drawn differing conclusions on whether S1P has a pro or anti-angiogenic role (Ben Shoham et al., 2012; Caballero et al., 2009; English et al., 2001; English et al., 2000; Gaengel et al., 2012; Licht et al., 2003). For lymphatic vessels, effect of S1P on patterning was described (Pham et al., 2010), and a matrigel implant *in vivo* model suggested an overall pro-lymphangiogenic effect (Yoon et al., 2008). While it appeared likely that S1PR1 represents the receptor transducing S1P effects in LECs similar to BECs, there was no clear demonstration of this from earlier studies.

This study shows, for the first time, that S1P-S1PR1 activity affects lymphangiogenesis by promoting VEGF-A–dependent LEC proliferation and by limiting LEC migratory and filopodia-forming responses *in vivo*. Filopodia formation and proliferation are hallmarks of endothelial tip and stalk cells, respectively. These cell phenotypes are key players in vascular patterning, and thus we suggest that the vascular network morphology is critically dependent on tightly controlled S1P-S1PR1 signaling.

While it was previously shown that S1PR1 signaling limits migratory, filopodia-forming responses during blood vessel development (Ben Shoham et al., 2012; Gaengel et al., 2012; Jung et al., 2012), a striking synergistic action of S1P with VEGF-A in stimulating cell proliferation, resulting in lymphatic vessel enlargement *in vivo*, was not yet shown and is surprising. As earlier studies demonstrated that VEGF-C/VEGFR-3 signaling is required for lymphangiogenesis during the first days after birth (Karpanen et al., 2006; Ochsenbein et al., 2016), we conclude that elevated S1P amplifies VEGF-A signaling in addition to ongoing VEGF-C growth factor signals. Therefore, under physiological conditions, VEGF-C signaling does not saturate LEC proliferation, which can be further enhanced by combined action of VEGF-A and S1P. In support of this, ectopic expression of VEGF-A was shown to increase proliferation rather than migration and filopodia-formation of LECs (Nagy et al., 2002; Wirzenius et al., 2007). Likewise, neutralization of VEGF-A alone in our developing diaphragm model, in the absence of enhanced S1P signaling, had no effect on assessed morphological parameters. This finding is in line with previously published results (Ochsenbein et al., 2016; Zhang et al., 2018), where genetic or antibody-mediated loss of VEGFR-2 signaling had no impact on postnatal lymphangio genesis.

Inhibition of S1PR1 signaling with the specific antagonist NIBR0213 led to chylus ascites formation and increased filopodia formation as well as branch formation *in vivo*, demonstrating dysfunctional and malformed lymphatic vessels. A similar phenotype was obtained in animals where S1PR1 was specifically deleted in LECs, emphasizing the important LEC-intrinsic signaling role of S1P/S1PR1. Of note, our data are well in line with a previous report showing increased filopodia formation in mice with lymphatic endothelial cell-specific depletion of sphingosine kinase, which is needed for the generation of S1P (Pham et al., 2010). Furthermore, as increased filopodia-formation is also seen in S1PR1-deficient blood vessels, our data establish a pan-endothelial mechanism.

For lymphatics we postulate that S1PR1-mediated inhibition of sprouting acts on alternative growth factor-driven lymphangiogenic processes, independent of VEGF-A. Earlier work showed that VEGF-C is a crucial driver of postnatal lymphangiogenesis (Zarkada et al., 2015; Zhang et al., 2018) and it will be interesting to investigate how S1P and VEGFC signaling converge *in vivo* in the future.

In order to gain better mechanistic insight into how S1P-S1PR1 affects VEGF-A-mediated lymphangiogenic processes we observed *in vivo*, we characterized S1P-S1PR1 signaling in LECs *in vitro*. We confirmed that S1P-S1PR1 signals in a constitutive and autocrine fashion, as expected from earlier genetic studies showing that LECs generate S1P in a sphingosine kinase 1/2 -dependent manner in mice (Pham et al., 2010). Surprisingly, our *in vitro* data clearly showed that VEGF-A can only signal and exert its mitogenic function when autocrine, constitutive S1P-S1PR1 signaling is active in LECs. The VEGF-A-mediated proliferative response of LECs was completely inhibited following blockade of S1PR1. In line with this finding, antagonizing S1PR1 also strongly inhibited VEGF-A – stimulated ERK1/2 activity. Interestingly, 50% of VEGF-A-induced ERK1/2 activity seems to be S1PR1-independent. However, the proliferation data demonstrate that S1PR1-independent ERK1/2 activation is not sufficient to translate into a functional output.

A previous report in HUVECs suggested a decrease of VEGF-A-induced pERK1/2 levels with S1P co-treatment (Gaengel et al., 2012). In our experience, HUVECs responded in a similar way to LECs by increasing the pERK1/2 levels. Taken together, we propose that S1P-S1PR1 signaling is a constitutive, LEC-intrinsic mechanism enhancing VEGF-A signaling and proliferative function. In line with this conclusion, sphingosine kinases were shown to be important for VEGF-A-mediated ERK1/2 activation in retinal blood endothelial cells (Maines et al., 2006) and in a bladder tumor cell line (Shu et al., 2002).

Upon VEGF-A stimulation, VEGFR-2 is internalized and traffics through different intracellular compartments, such as Rab5+ endosomes (Simons et al., 2016). These steps are important for VEGF-A-mediated downstream signaling and aberrant trafficking of VEGFR-2 was shown to decrease ERK1/2 activation (Lanahan et al., 2010). As deficient S1PR1 signaling decreased VEGFR-2 trafficking to the perinuclear region, we speculate that S1PR1–mediated VEGFR-2 trafficking is needed for efficient ERK1/2 activation. Furthermore, as S1PR1 antagonism does not impact VEGFR-2 trafficking to Rab5+ early endosomal compartments, we conclude that the crosstalk between VEGFR-2 and S1PR1 signaling happens between early endosomes and compartments in the perinuclear region. Interestingly, others have shown that S1PR1 and VEGFR-2 can form signaling complexes (Bergelin et al., 2010), and both VEGFR-2 and S1PR1 have been shown to traffic through the same (Rab5+) endosomal compartments (Kofler et al., 2018; Martinez-Morales et al., 2018) relevant for ERK1/2 signaling (Kofler et al., 2018). Furthermore, sphingosine kinases play an important role in endocytotic membrane trafficking in general (Lima et al., 2017; Shen et al., 2014).

Interestingly, similar hyperproliferative lymphatic vessel phenotypes as the ones induced by enhanced S1P signaling in our study were described in mice with deficient negative regulators of ERK1/2 signaling, namely SPRED1 and RASA1, as well as for gain-of-function of RAF1, a positive regulator of ERK1/2 (Deng et al., 2013; Lapinski et al., 2012; Taniguchi et al., 2007). Further, lymphatic-specific deletion of the transcription factors FOXC1 and FOXC2 decreased *Rasa3* in embryonic LECs, increased *S1pr1* levels and activated ERK1/2 that resulted in lymphatic vessel enlargement without changes in branching (Fatima et al., 2016). Considering these findings together with our finding that S1P-S1PR1 controls VEGF-A-induced ERK1/2 activation, we suggest that S1P acts as an enhancer of the Ras-Raf-MEK-ERK1/2 signaling cascade upon VEGF-A stimulation *in vivo*, and that negative modulation of S1P-S1PR1 signaling may be beneficial in treating certain genetic diseases linked to lymphatic hyperproliferation.

In summary, we show for the first time that autocrine S1P-S1PR1 influences lymphangiogenesis by promoting VEGF-A-dependent LEC proliferation and by limiting migratory and filopodia-forming responses. By doing so, S1P-S1PR1 signaling may favor generation of proliferative stalk cells over filopodia-forming tip cells. In line with this notion, S1PR1 negative blood endothelial cells preferably acquire a tip cell position in the growing retinal vasculature (Gaengel et al., 2012). The novel mechanistic insight into the modulation of VEGF-A-mediated lymphangiogenesis provided here could be relevant for the understanding of diseases characterized by prominent hyperplastic lymphatic vessels and loss of lymphatic function such as RASopathies.

## Author contributions

A.M.G.O., G.J. and K.S. designed experiments and A.M.G.O. wrote the paper with support of G.J. and K.S.; G.J. and K.S. supervised the study; A.M.G.O., B.W., S.V. and C.B. performed experiments and A.M.G.O. analyzed data; C.G., L.C., S.L. supported performance of experiments and experimental design; N.H. wrote the personalized Fiji macro and helped with picture processing.

## Acknowledgements

The authors thank Samuel Barbieri, Dominic Trojer, Isabelle Claerr and the Novartis lab animal services for their support with experimental procedures, Dr. Andreas Billich, Dr. Frederic Bassilana, Dr. Marc Bigaud, Dr. Laure Bouchez and Prof. Michael Detmar for scientific input, Dr. Anne Granger for critical reading of the manuscript and the Novartis postdoc program and Dr. Tewis Bouwmeester for their guidance.

