## Supplementary data for "S1P-S1PR1 activity controls VEGF-A signaling during lymphatic vessel development"

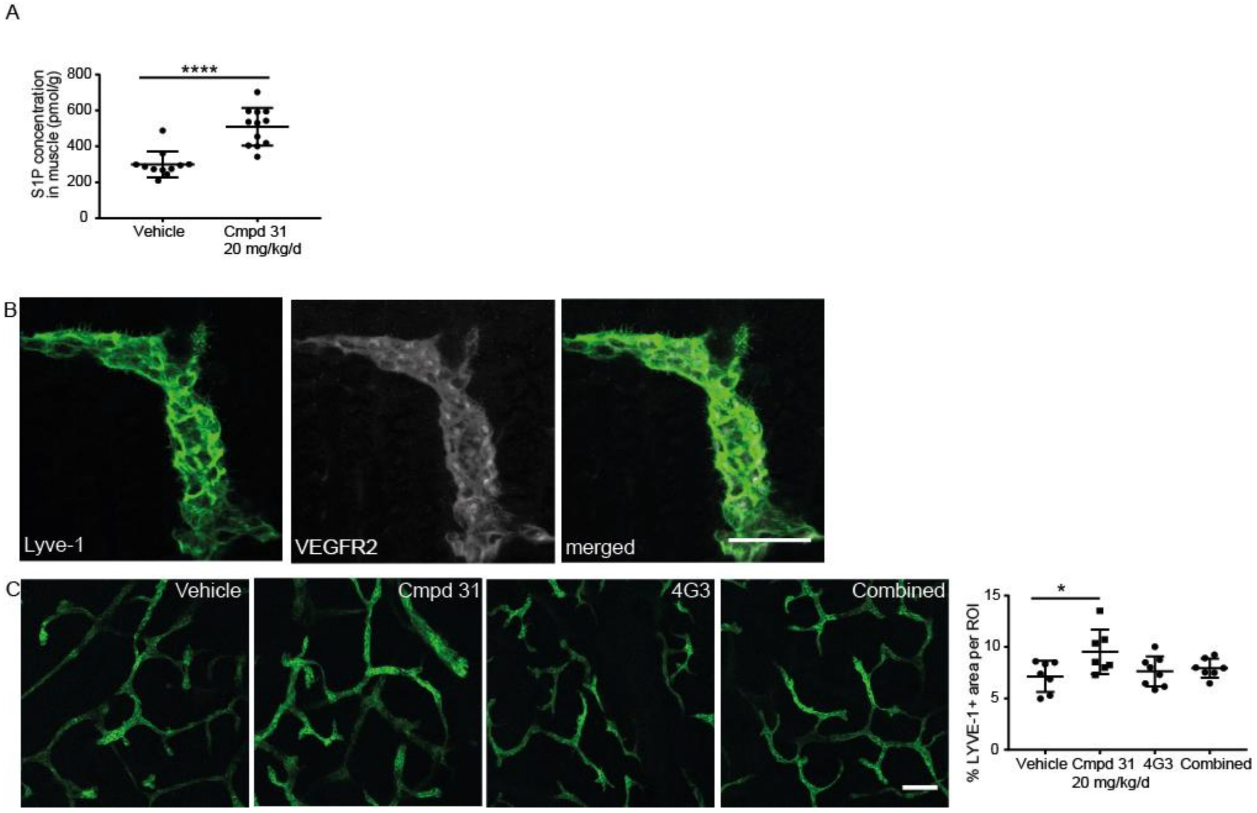


**Fig S1: Effects of S1P on lymphatic vessels in the diaphragm and on lymphangiogenic processes in skin.** A) Quantifications of S1P concentration of vehicle and Cmpd 31 treated pups. B) Representative confocal images of P2.5 diaphragm whole mount stained for LYVE1 (green) and VEGFR-2 (grey), scale bar 50 μm. C) Confocal images and quantification of LYVE1 immunostained abdominal skin whole mount of P4.5 pups that received indicated treatments, scale bar 100 μm. Statistical analyses: Dots represent data from individual animals, lines represent the group means, error bars = ±SD. Statistical significance was calculated with an unpaired Student's t-test (A) and with one-way ANOVA and Dunnett's test (C).

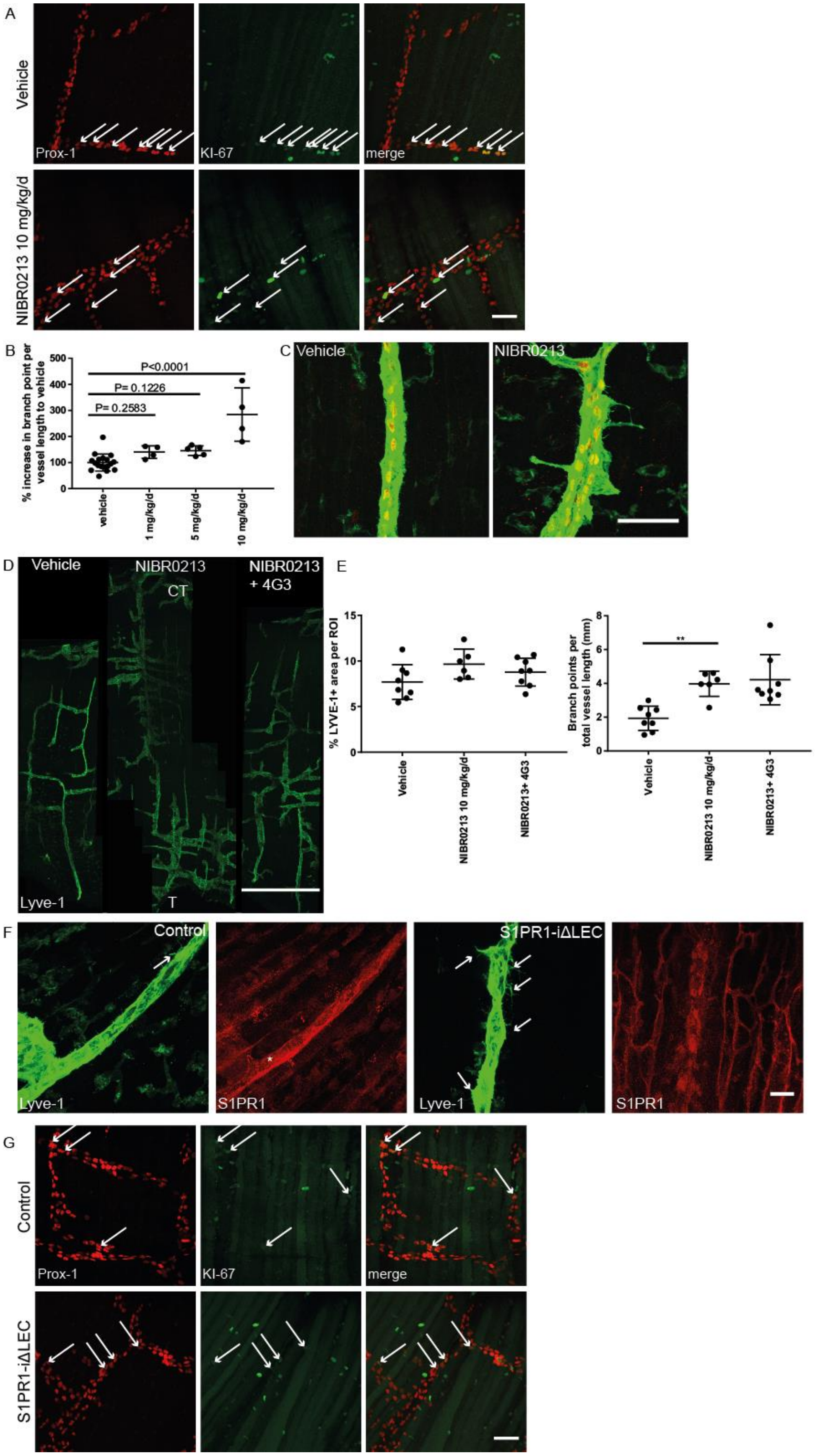
A
Vehicle
NIBR0213 10 mg/kg/d
Prox-1
KI-67
merge
B
% increase in branch point per vessel length to vehicle
P<0.0001
P= 0.1226
P= 0.2583
vehicle
1 mg/kg/d
5 mg/kg/d
10 mg/kg/d
C
Vehicle
NIBR0213
D
Vehicle
NIBR0213
CT
NIBR0213 + 4G3
Lyve-1
T
E
% LYVE-1+ area per ROI
Branch points per total vessel length (mm)
**
Vehicle
NIBR0213 10 mg/kg/d
NIBR0213+ 4G3
F
Control
S1PR1-iΔLEC
Lyve-1
S1PR1
G
Control
S1PR1-iΔLEC
Prox-1
KI-67
merge

**Fig S2: S1PR1 inhibition has no effect on LEC proliferation while it increases lymphatic vessel branching and filopodia formation in a VEGF-A-independent manner. S1PR1 expression is reduced specifically on the lymphatic vessels in S1PR1-iΔLEC pups.** A) Representative confocal images of diaphragm whole mounts immunostained for PROX-1 (red) and KI-67 (green) after indicated treatments. Arrows: KI-67 + LECs, scale bar 50 μm. B) Quantification of lateral segments of LYVE1 immunostained diaphragm whole mounts of vehicle or 1, 5 or 10 mg/kg/day NIBR0213 treated pups. C) High magnification confocal images of diaphragm whole mounts immunostained for PROX-1 (red) and LYVE1 (green) focusing on filopodia after indicated treatments, scale bar 50 μm. D) Representative confocal images of P4.5 diaphragm whole mounts (pleural side) stained for LYVE1 of vehicle, 10 mg/kg/day NIBR0213, or NIBR0213 and 6 mg/kg/day 4G3 treated pups, scale bar 1 mm. E) Quantification of lymphatic vessel parameters of treated pups. F) Confocal images of S1PR1 (red) and LYVE1 (green) immunostained diaphragm whole mounts of P7 control or S1PR1-iΔLEC pups. Arrows point at filopodia, Scale bar 20 μm. G) Representative confocal images of diaphragm whole mounts immunostained for PROX-1 (red) and KI-67 (green), comparing P7 control or S1PR1-iΔLEC pups. Arrows: KI-67 + LECs, scale bar 50 μm. Statistical analyses: Dots represent data from individual animals and group means ± SD are shown. Statistical significance was calculated using one-way ANOVA and Dunnett’s test (B) and Kruskal-Wallis and Dunn’s test (E).

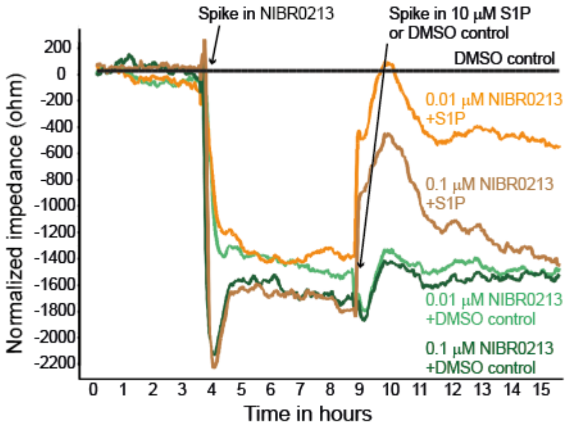


**Fig S3: NIBR0213 is not toxic for LEC.** Normalized (to DMSO control) electrical impedance of LEC monolayers after indicated treatments. Excess S1P competes with NIBR0213 for S1PR1 receptor binding and induces barrier tightening.

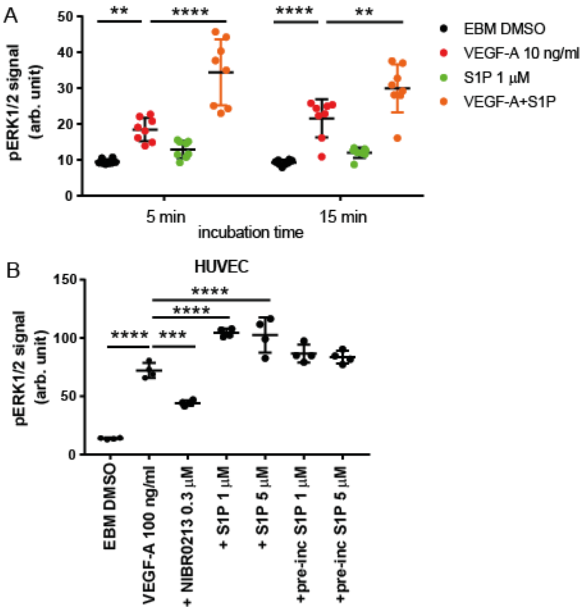


**Fig S4: S1P modulation affects growth factor-induced downstream signaling.** A) Quantification of pERK1/2 levels in LEC lysates after indicated treatments. B) Analysis of pERK1/2 levels assessed by HTRF in HUVEC lysates after 10 min of indicated treatments. Statistical analyses: Dots represent individual measurements, lines represent group means. Error bars = ±SD. Statistical significance was calculated using ANOVA and Dunnett's test.

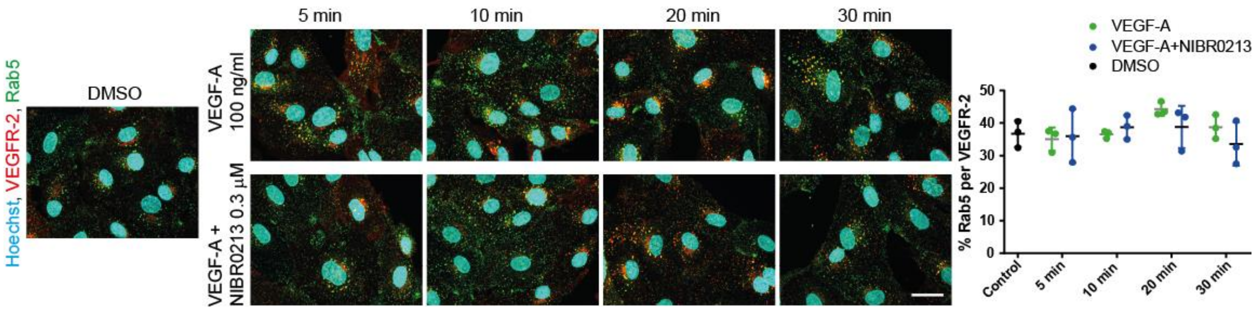


**Fig S5: S1PR1 activity does not influence VEGFR-2 co-localization with Rab5 after VEGF-A treatment.** Immunofluorescent staining of VEGFR-2 (red), Rab5 (green) and Hoechst (blue) and quantification of Rab5 and VEGFR-2 co-localization in cultured LEC after indicated treatments and incubation times, scale bar: 50 µm. Statistical analyses: Dots represent individual measurements, lines represent group means, error bars = ±SD. Statistical significance was calculated with a Student's t-test (compared to VEGF-A at corresponding time point).
